# Unilateral online ultrasound stimulation of early visual cortex suppresses responses to contralateral visual stimuli

**DOI:** 10.1101/2025.08.05.668165

**Authors:** Suraya Dunsford, Keith Murphy, Ema Darrieutort, Elsa Fouragnan, Giorgio Ganis

**Affiliations:** School of Psychology, Faculty of Health, University of Plymouth, Plymouth, United Kingdom; Brain Research and Imaging Centre, Faculty of Health, University of Plymouth, Plymouth United Kingdom; Stanford University, California, United States; Attune Neurosciences, California, United States

**Author notes:** These authors contributed equally to this work. Primary corresponding author: Suraya Dunsford, Additional corresponding authors: Giorgio Ganis, Elsa Fouragnan.

**Keywords:** Transcranial ultrasound stimulation, visual cortex, real-time, visual evoked potentials, online, non-invasive brain stimulation, neuromodulation

## Abstract

Transcranial ultrasound stimulation (TUS) shows great promise for inducing neuroplastic changes that persist long after stimulation. Evidence of stimulation-locked neural changes would enable closed-loop application of TUS, but such responses have not yet been clearly dissociated from the coincident neural response to auditory and peripheral stimulation associated with TUS. We leveraged the contralateral retinotopic organization of the early visual cortex to isolate online TUS effects from peripheral confounds in 19 subjects. Using a hemifield visual stimulation paradigm combined with high-precision, functional MRI guided TUS, we applied TUS to the left early visual cortex while participants viewed checkerboards presented in the left or right visual field. TUS was delivered randomly on half of the trials, enabling within-subject comparisons of pattern-locked visual evoked potentials (VEPs) across hemispheres and against no stimuli. We observed a reduction in VEPs in the contralateral, but not ipsilateral, hemifield, consistent with genuine online neuromodulatory effects. Furthermore, online suppression was positively correlated with the TUS dose delivered to the target, as estimated by modelling TUS field-target overlap and differential attenuation through heterogeneous skulls. Collectively, these findings provide a robust framework for future studies aiming to map the TUS parameter space in real time by leveraging topographic organization to control for peripheral confounds.

## Introduction

Transcranial ultrasound stimulation (TUS) is a non-invasive neuromodulation technique offering precise access to the deep brain without requiring any surgical implantation (1, 2). By delivering focused acoustic energy through the skull, TUS can achieve millimeter-scale precision, making it particularly well-suited for modulating and studying discrete neural circuits (3). Importantly, the low form factor afforded by TUS makes it particularly well suited for real-time closed loop modulation during tasks. However, direct evidence of real time (online) effects during stimulation has remained elusive in human research.

Although TUS has demonstrated the ability to modulate brain activity, a key distinction exists between its offline and online effects (4). Effects that persist after stimulation, also known as offline effects, are thought to reflect neuroplastic changes (5, 6). These are especially promising for clinical applications targeting long-term functional changes and symptom relief, without the need for constant stimulation (7). In stark contrast, effects which occur during stimulation, or online effects, remain almost entirely unexplored. Suprathreshold effects, including direct action potential induction and behavioral responses, have been demonstrated in animal models (8-10) but have not yet been reliably demonstrated in humans. This discrepancy may reflect a combination of multiple factors, such as greater ultrasound attenuation and scattering by the human skull, reduced targeting precision, or limited stimulation intensity due to safety constraints in humans. As a result, most human TUS studies to date have reported subthreshold modulation of neural activity, without clear evidence of direct evoked neural effects (11, 12). Nevertheless, online effects offer a critical opportunity to rapidly interrogate the causal relationship between TUS parameters and neural circuit activity (11). They also offer the unparalleled opportunity to support real-time closed-loop neuromodulation paradigms which is particularly well suited for TUS, given the lightweight nature of ultrasound transducers (13). However, auditory, bone-conducted sound, and other peripheral nerve stimulation artifacts can occur with TUS, and independently evoke neural and behavioral responses (14, 15). Thus, isolating true online effects from these effects is particularly challenging. Although techniques such as sham stimulation, auditory masking (16), and careful parameter tuning are increasingly employed to minimize these confounds (13), residual perceptual artifacts often persist. Consequently, progress in leveraging online TUS effects will depend critically on rigorous experimental designs capable of dissociating true neuromodulatory effects from confounding sensory-driven responses.

The early visual cortex, comprising V1 within the calcarine sulcus and neighboring V2 (17), serves as an ideal testbed for studying neuromodulation techniques due to its well-defined contralateral retinotopic organization (18-20). By presenting hemifield-specific visual stimuli, it is possible to assess the spatial specificity of neuromodulation effects, based on the contralateral organization of the early visual cortex. Spatial specificity can then be used to rule out peripheral confounds which should not show hemispheric lateralization. Demonstrations of this logic have been provided using transcranial magnetic stimulation (TMS). For example, single TMS pulses delivered to the occipital pole in one hemisphere can cause phosphenes and, at higher intensities, transient scotomas in the contralateral visual hemifield (e.g., 21, 22). TMS to early visual cortex has also been shown to affect visual evoked potentials (VEPs) in a spatially-specific manner. Thut and colleagues (23) found that the topography of the visual P1 and N1 components elicited by a full-field checkerboard was modulated by single pulse TMS stimulation of the occipital pole. Furthermore, Reichenbach and colleagues (24) showed lateralized effects on the visual P1 component elicited by a lower right quadrant checkerboard following single pulse TMS stimulation of the occipital cortex. Collectively, these studies illustrate the utility of using the contralateral retinotopic organization of early visual cortex as a powerful approach for assessing the online effects of neuromodulatory techniques.

To extend this spatially specific approach to TUS we envisioned hemifield-specific visual stimuli, where researchers can elicit lateralized responses in left and right early visual cortex, providing an internal control for spatial specificity. This approach is particularly useful to differentiate genuine neuromodulatory effects from auditory confounds, which should produce more global and bilateral cortical activation (14, 25). If TUS effects are truly visual, they should selectively modulate neural activity in the hemisphere contralateral to the stimulated visual field, whereas non-specific artifacts are more likely to affect activity in both hemispheres. Thus, neuromodulation of the early visual cortex not only enables precise temporal tracking of online effects but also offers a robust framework for disentangling them from pervasive confounding signals.

Prior work by Nandi and colleagues (26) applied online TUS to the occipital cortex during full-field visual stimulation and observed modulation of the early N75 VEP component, suggesting that online TUS can affect early visual processing. Related studies (e.g., 27) have also reported TUS-induced modulation of visual responses, though limited detail regarding lateralization in their design constrains interpretability in relation to the current study. The present study introduces three key methodological refinements to improve spatial specificity and better control for confounding factors.

First, we used hemifield-specific pattern-onset stimuli to elicit clear contralateral VEPs (e.g., 28). Leveraging the contralateral organization of early visual cortex, we compared TUS effects during right versus left hemifield stimulation, while TUS was always applied to the left early visual cortex on 50% of trials in a fixed alternating sequence. This design allowed for a direct comparison between TUS effects during right hemifield stimulation (processed by the targeted left visual cortex) and left hemifield stimulation (processed by the non-targeted right visual cortex, serving as a within-subject control). Critically, this setup controlled for potential auditory confounds, as any non-specific auditory effects of TUS would be present in both trial types and should cancel out in differential analyses. We used an 8 second interval between consecutive TUS trials, separated by a sham trial, to ensure tissue temperature rise remained within ITRUSST safety guidelines.

Next, we chose TUS parameters which would maximize acoustic energy deposition while minimising auditory artifacts. Previous work in animal studies from our team have shown that lower pulse repetition frequency, higher duty cycle pulses produced a faster, higher amplitude calcium response across several brain areas (10). At close inspection, elicited calcium rise does not peak for at least several hundred milliseconds, suggesting that a single longer pulse would elicit the largest instantaneous effect; coincident with findings in culture (29). Fewer pulses will also reduce the audible clicking sound which occurs at the edge of singular pulses and can produce confound to studies (25). Taking these findings, our experimental paradigm timing, and thermal and mechanical safety into account, we designed a schedule where a single 500 ms pulse was initiated 400 ms prior to and entirely overlapping a 100 ms visual stimuli, maximizing the expected overlap of neuromodulatory effect with VEPs.

Finally, we employed individual fMRI-guided neuronavigation to target activated portions of the left early visual cortex within the calcarine sulcus. The core logic of this design is that if TUS modulates neural activity in the targeted region, it should selectively alter early VEPs following right, but not left, hemifield stimulation (e.g., 28). We hypothesised that online TUS would modulate early visual processing, producing measurable changes in VEPs within 100 ms post-stimulus, particularly at left posterior electrodes closest to the targeted region.

## Materials and Methods

### Participants

Twenty healthy volunteers (14 female, 20 - 60 years old, mean = 32.5, SD ± 12.9), participated in the study and were recruited via online advertising and posters distributed in the local community. An online survey followed by telephone interviews was used to screen all participants to ensure inclusion of healthy participants without any vision impairments, aged between 20–60 years with fluency in spoken English. One participant was excluded from the analyses due to an intermittent equipment issue that compromised the EEG recordings, leaving nineteen participants (14 females, 20 – 60 years old, mean = 33.1, SD ± 13.0) for the analyses.

Exclusion criteria, specifically for TUS and MRI safety, included: 1) self-reported pregnancy, 2) psychoactive drug use or current use of any medication affecting the central nervous system or seizure threshold (unless deemed safe and non-interfering), 3) a first-degree relative with a history of epilepsy, convulsions or seizures, 4) extreme mood fluctuations, 5) current psychiatric diagnosis or medication, 6) any history of neurological illness or disorder, or moderate-severe brain injury, 7) any additional known contraindications for MRI scanning.

The University of Plymouth Faculty of Health Staff Research Ethics and Integrity Committee approved the study (reference ID: 2021-2398-1498; date: 18/01/2021). Before participation, all individuals provided written informed consent after receiving a full explanation of the experimental procedures. Participants were compensated £15 per hour spent in the lab, and reimbursed up to £10 per session for travel expenses. All study sessions were conducted at the Brain Research and Imaging Centre in Plymouth, United Kingdom.

### Experimental Design

Figure 1 summarizes the study design and procedures. Participants underwent an initial MRI session to obtain a high-resolution T1-weighted image, a Pointwise Encoding Time Reduction with Radial Acquisition (PETRA) image for skull information for acoustic and thermal simulations, and functional visual localizers for target location. The visual localizers included blocks to map the visual hemifields and the lower quadrants. Using these data, a left hemisphere stimulation target was identified and the stimulation trajectory was optimized before the second session using K-Plan, a GUI executing k-wave simulations (30) to optimise targeting and ensure safety. Specifically, the stimulation target was selected to meet the following criteria: (i) it included a cluster of voxels in the left calcarine sulcus that were activated by right hemifield stimuli, (ii) it was at least 28 mm distant from the scalp, to reduce the risk of excessive soft tissue and skull heating; note that this depth requirement placed the target in regions representing relatively peripheral regions of the visual field, and (iii) it was at least 10 mm away from the edge of the right hemisphere to ensure that, even if the targeting system shifted slightly over time, the stimulation would remain confined to the left visual cortex.

**Fig. 1.**
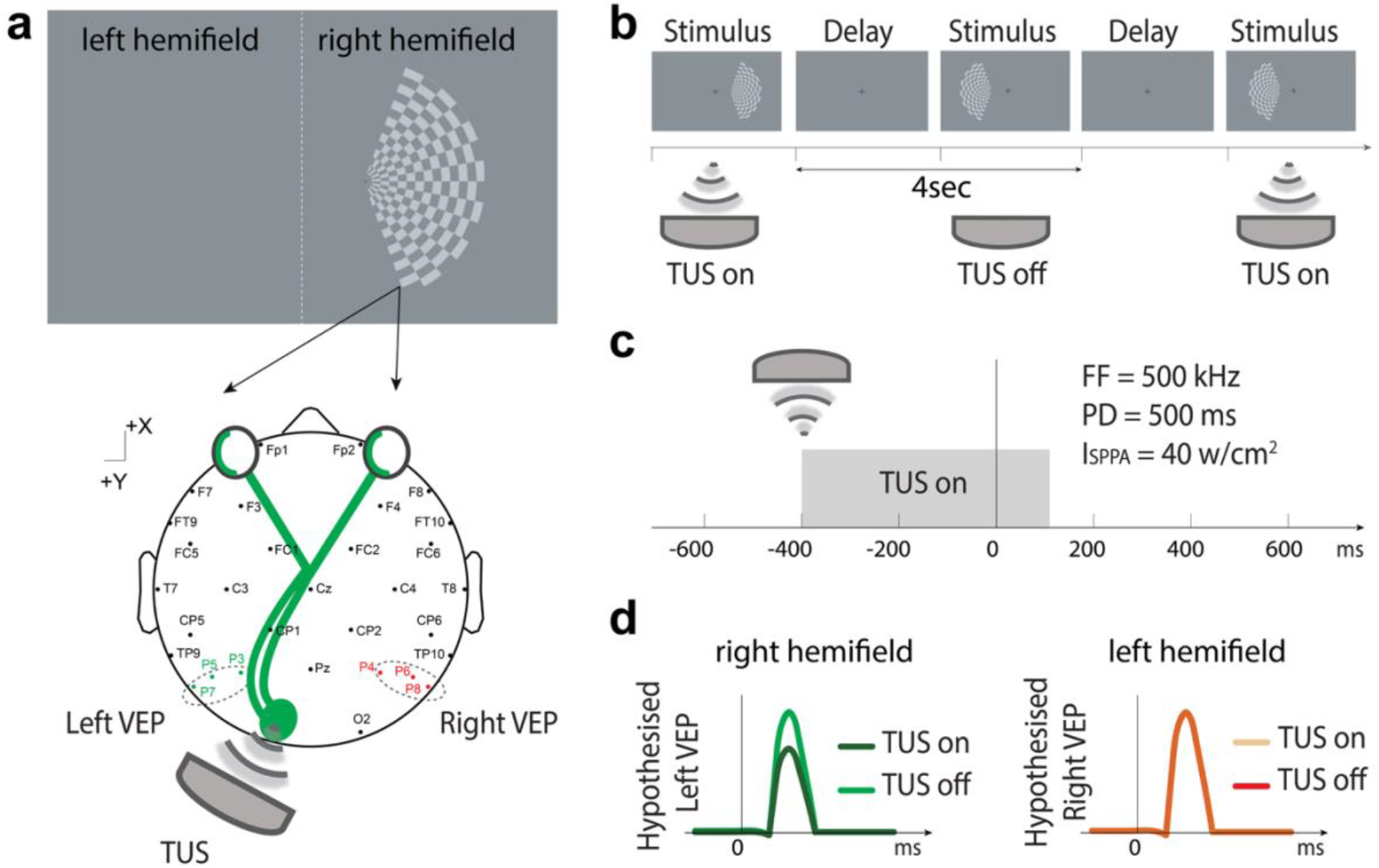
Experimental paradigm and hypothesized VEP modulation by TUS. **(A)** Schematic representation of visual stimulation, TUS and EEG recording. A checkerboard stimulus was presented in either the left or right visual hemifield while TUS was applied to the left early visual cortex. Stimuli presented in one hemifield were hypothesized to be processed in the contralateral visual cortex, and corresponding VEPs were recorded from bilateral parietal electrodes (green and red dots). TUS parameters are displayed but more detailed in the methods section. **(B)** Trial structure illustrating alternating trials of TUS on and off. A small fixation cross was present throughout the session. Each trial began with a briefly flashed visual stimulus (20 ms), followed by a delay. **(C)** Temporal alignment of the TUS waveform relative to the visual stimulus onset. TUS began 400 ms before visual stimulus onset and ended 80 ms after stimulus offset. **(D)** Hypothesized VEP modulation by TUS. **Left:** Right hemifield stimulation elicits left VEPs, predicted to be altered under TUS. **Right:** Left hemifield stimulation elicits right VEPs, which are not expected to be altered by TUS.

These steps informed the setup for the TUS session, which was planned and executed using neuronavigation (Brainsight, Rogue Research Inc.) to align the sonication focus with the individually defined target. MRI sessions were followed by online TUS sessions at least one week apart, during which TUS was applied to the target during the visual task.

### Magnetic Resonance Acquisition

Participants underwent a series of MRI scans on a Siemens MAGNETOM Prisma 3 T scanner (VE11E, Siemens Healthineers, Erlangen, Germany) with a 64-channel head coil. The sequence of scans was as follows:

1. Initial scout scan for accurate localization of the participant
2. T1-weighted sequence, image acquisition parameters were: 2100.0 ms repetition time (TR), 2.26 ms echo time (TE), 900 ms inversion time (TI), and a flip angle of 8°. Imaging was performed over a 256×256 mm^2^ field of view with a base resolution of 256, yielding isotropic voxels of 1.0×1.0×1.0 mm^3^. The acquisition comprised 176 slices per slab with 18.2% slice oversampling, and phase partial Fourier was set to 6/8. Parallel imaging was applied using GRAPPA with an acceleration factor of 2 (24 reference lines), and no additional 3D acceleration was used. Key sequence parameters include an echo spacing of 6.8 ms, a turbo factor of 208, and a bandwidth of 200 Hz/Px.
3. fMRI task, image acquisition parameters were: TR = 2000 ms, TE = 30.00 ms, and a flip angle of 72°. The acquisition employed a voxel size of 2.0×2.0×2.0 mm^3^ across a 216×216 mm^2^ field of view, with 60 slices (slice thickness = 2.00 mm) covering the occipital cortex, and oriented approximately orthogonally to the calcarine sulcus. A total of 190 volumes were collected. Parallel imaging was implemented using GRAPPA with an acceleration factor of 2 (24 reference lines, FLEET reference scan mode), in combination with a multi-band acceleration factor of 2 to enhance temporal resolution. Further, the EPI sequence featured an echo spacing of 0.64 ms, an EPI factor of 108, and a bandwidth of 2314 Hz/Px to optimise image quality and reduce distortions.
4. Field map (all measurement parameters made to match fMRI acquisition as far as possible).
5. PETRA (Pointwise Encoding Time Reduction with Radial Acquisition) image acquisition parameters were: TR = 3.61 ms, TE = 0.07 ms, and a flip angle of 1.0°. The scan was acquired with a 240 × 240 mm^2^ field of view at a base resolution of 320, yielding an in-plane resolution of approximately 0.75 mm with 0.75-mm slice thickness (resulting in reconstructed voxels of approximately 0.8×0.8×0.8 mm^3^). A total of 320 slices per slab were acquired utilising 60000 radial views. Additional sequence optimizations included a bandwidth of 359 Hz/Px, active acoustic noise reduction, and a “Whisper” gradient mode.

### EEG/MRI Stimuli

During the EEG/TUS session, visual stimuli consisted of radial half-checkerboards presented in random order to either the left or right visual hemifield on separate trials (Fig. 1A). The diameter of the checkerboards subtended about 20 degrees of visual angle. The screen background was set to grey (RGB: 128,128,128), and Michelson contrast was set to 25%. Stimuli were displayed for 20 ms (2 frames) using a 24 inch monitor with refresh rate of 100 Hz. Similar stimuli were used for the MRI localizers. However, during MRI acquisition, the checkerboards were counterphase-flickered at 8.5 Hz to maximize visual cortex engagement. In the scanner, stimuli were presented using a Cambridge Research BOLDScreen 32 (1920 × 1080, refreshed at 60 Hz) placed behind the bore and seen by the participants via a mirror mounted on the head coil.

### EEG Acquisition

EEG data was acquired at 500 Hz from 32 active electrodes (actiCAP slim) with an ActiCHamp PLUS amplifier (Brain Products) using BrainVision Recorder. The electrodes were mounted with actiCAP snap electrode holders on a cap modified by cutting a circular opening over the left occipital cortex to accommodate the ultrasound transducer. As a result, O1 and Oz sites could not be used, and their corresponding amplifier channels were instead reassigned to record data from the additional sites P5 and P6. This resulted in the following electrode configuration: O2, P7, P5, P3, Pz, P4, P6, P8, TP9, CP5, CP1, CP2, CP6, TP10, T7, C3, Cz, C4, T8, FT9, FC5, FC1, FC2, FC6, FT10, F7, F3, Fz (reference), F4, F8, FP1, Fpz (ground), and FP2.

### Ultrasound Stimulation

We used the NeuroFUS TPO system (Brainbox Ltd., Cardiff, UK) with a 500 kHz central frequency transducer. The transducer had a 64 mm diameter and a four-channel annular array with four fundamental mode impedance matching networks. Online TUS stimulation was delivered trial-wise, with each trial consisting of a 500 ms pulse of 500 kHz continuous-wave ultrasound at an ISPPA of approximately 40 W/cm^2^. The stimulation onset was time-locked to occur 400 ms prior to the onset of the visual stimulus. Each block comprised 100 trials (50 stimulation, 50 sham, alternating) with an inter-trial interval of 4.0 sec. A total of 5 blocks were administered per session.

The spatial peak pulse average intensity (ISPPA) in water was kept consistent at ∼40 W/cm^2^ for all participants. These values were validated using hydrophone tank measurements (Yaakub et al., 2023b). Stimulation protocols conformed to the human safety guidelines proposed by the ITRUSST consortium (31). Acoustic simulations predicted a maximum skull temperature rise of <2°C for the majority of participants. We also computed the Cumulative Equivalent Minutes at 43°C (CEM43), a conservative metric quantifying thermal exposure relative to the threshold for cellular damage. In all participants, CEM43 values remained well below 0.25, and in our dataset, never exceeded 0.1 (see Supplementary Table 1). Similarly, the transcranial Mechanical index never exceeded 1, far below the 1.9 advised by ITRUSST consortium (31).

**Table 1.** Ultrasound parameters across different target tissues for left early visual cortex.

| Tissue | Thermal Dose<br>(CEM43°C) |  | Pressure (MPa) |  | ISPPA<br>(W/cm<sup>2</sup>) |  | Mechanical Index |  |
| --- | --- | --- | --- | --- | --- | --- | --- | --- |
|  | *M* | *SD* | *M* | *SD* | *M* | *SD* | *M* | *SD* |
| ROI mask | 0.00 | 0.00 | 0.31 | 0.09 | 3.90 | 1.72 | 0.44 | 0.13 |
| Soft tissue | 0.01 | 0.01 | 0.39 | 0.07 | 5.28 | 1.60 | 0.55 | 0.10 |
| Skull | 0.03 | 0.03 | 0.52 | 0.13 | 9.30 | 4.74 | 0.73 | 0.19 |

### Experimental Procedure

Following EEG cap placement, the participant’s hair was prepared to facilitate effective coupling. The ultrasound transducer was coupled to the scalp using an aqueous ultrasound gel (Aquasonic 100, Parker Laboratories Inc.) and a gel pad (Aquaflex, Parker Laboratories Inc.), following the procedure outlined by Murphy et al. (13). This method ensured minimal air entrapment between the transducer face and the scalp, reducing acoustic reflection and preserving optimal acoustic energy transmission to the target region (32).

Subsequently, participant registration was conducted using BrainSight (Rogue Research Inc.). Participants were seated in a chair with their chin resting on a chin rest. Two microfiber-padded positioning arms were gently placed against the zygomatic bones to maximize comfort and maintain head stability throughout the procedure. The ultrasound transducer was then aligned and securely mounted using the arm of the DuoMag system (Deymed Diagnostic s.r.o.; https://deymed.com/duomag-xt), fitted with a custom 3D-printed adapter designed to hold both the transducer and the tracking sensor.

#### Visual task

Participants sat in a dimly illuminated room (∼15 lx), 57 cm from the screen, maintained central fixation throughout the task, and used their dominant hand to press the left or right key in response to checkerboards presented in the left or right visual field, respectively. The task was split into 5 blocks (∼410 seconds each), each with 100 trials, divided equally between the 2 × 2 combination of condition (TUS, Sham) and visual hemifield (Left, Right) (Fig. 1B). Each block contained 25 trials of each type, pseudo-randomly distributed. At the end of each block there was a 10 sec rest period. These rest periods were also used to pause the study and adjust TUS transducer position, if the focus had drifted away from the target during the preceding block. White noise (∼55 dB) was used throughout the session, to mask any potential TUS-elicited sounds participants might have heard. At the end of the session, when asked, five participants reported hearing a sound at some point during the session that may have been related to the stimulation.

### Statistical Analysis

#### Behavior

Accuracy data were analysed with a 2 × 2 repeated measures ANOVA, with condition and hemifield as factors, to ensure participants were paying attention to the stimuli. Note that response times were not analyzed as the task was intentionally unspeeded to discourage movement and minimize motion artifacts.

#### EEG

The EEG preprocessing pipeline includes activity recorded from 31 EEG sensors referenced to Fz and consists of the following preprocessing steps carried out in EEGLAB (33) and ERPLAB (34): (i) trimming of EEG data segments without event codes for more than 4 seconds; (ii) removal of outlier channels (excluding eye channels and anterior frontal channels to avoid removing EOG artifacts to be corrected with ICA later) due to paroxysmal artifactual activity, using a maximum variance threshold; (iii) high-pass filtering with a noncausal Butterworth filter (0.3 Hz half-amplitude cutoff, 12dB/oct roll-off, order = 2); (iv) low-pass filtering with a noncausal Butterworth filter (40 Hz half-amplitude cutoff, 12dB/oct roll-off, order = 2); (v) detrending (35), with *removeTrend* and a ‘detrendcutoff’ value of 0.2 Hz in the PREP pipeline (36) (vi) Removal of residual line noise, if still present after low-passing, (50Hz and the first 4 harmonics) using *CleanLineNoise* from the PREP pipeline, with default parameters (36); (vii) Removal of EEG data segments containing data at anterior frontal channels with absolute amplitude larger than 300 μV or peak-to-peak amplitude larger than 600 μV, using a 500 ms window shifted in 250 ms steps; (viii) Removal of EEG data segments at any other channels (other than anterior frontal) with absolute amplitude larger than 225 μV or peak-to-peak amplitude larger than 300 μV, using a 500 ms window shifted in 250 ms steps; (ix) removal of ocular or muscle artifacts using Independent Component Analysis (ICA, infomax and ICLabel (37) with a probability of being a muscle, heart, or channel noise artifact greater than 0.9; for ocular artifacts, removal of components for which the eye artifact weight is the largest and for which there is less than 5% brain activity weight; (x) Epoching between −200 and 800 ms for the primary analyses, and between −600 and 800 ms for the pre-stimulus analyses; (xi) interpolation of removed channels (if any); (xii) removal of any residual trials with amplitude at any channels larger than 75 μV in absolute value, using a 200 ms window shifted in 100 ms steps; (xiii) removal of any residual trials with peak-to-peak amplitude at channels P3, P4, P5, P6, P7, P8 larger than 100 μV, within a moving 200 ms window shifted in 100 ms steps.

Artifact-free epochs were averaged for each participant and condition to generate the VEPs. A 200 ms baseline was applied: before stimulus onset (−200 to 0 ms) for the primary VEP measurements, and before TUS onset (−600 to – 400 ms) for assessing the effects of TUS before visual stimulus onset.

The first analysis compared the amplitude of TUS and sham VEPs elicited by right hemifield stimuli. For this analysis, signals from the left parietal electrodes (P3, P5, and P7) were averaged together to form a single virtual electrode (VPL), providing a measure of activity closest to the site of TUS (Fig. 3B). This analysis focused on the first 160 ms post-stimulus, capturing the initial volley reaching V1 (∼60 ms), and its subsequent propagation to extrastriate visual cortical areas and beyond (e.g., *38, 39*). To assess significance, eight sequential paired t-tests were conducted across 20 ms windows (0–160 ms). The symmetric right parietal electrodes (P4, P6, and P8) were similarly averaged to form a right-hemisphere control site (VPR), and the same analysis was repeated for VPR, to confirm that there were no differences between TUS and sham in the control hemisphere. Critically, we then used the same windows to compare the TUS–sham difference between VPL and VPR, calculated as (LTUS-LSham) - (RTUS-RSham). This approach allowed us to isolate “pure” TUS effects that were not contaminated by sound-related artifacts, as any such artifacts would be present in both hemispheres.

An additional analysis examined TUS-induced effects prior to visual stimulus onset (−400 to 0 ms). Here, the differences between TUS and sham conditions were tested across eight sequential 50 ms windows for both the VPL and VPR site averages.

For the amplitude analyses across multiple time windows, we controlled for multiple comparisons using the MATLAB implementation of the Benjamini–Hochberg method (40).

Finally, we computed the 50% fractional peak latency for the P1 component, using function *pop_geterpvalues* in ERPLAB.

#### MRI

Unfolded cortical surfaces were automatically derived from the T1 datasets using *recon-all* in FreeSurfer (41) and were used to display cortical curvature maps in cases where identification of parts of the calcarine sulcus was ambiguous. FMRI data were analyzed using AFNI v21.1.20 (42) and SPM12. No spatial smoothing was applied to the functional timeseries. Motion-correction was carried out using function *3dvolreg* (AFNI), aligning all volumes to a reference at the beginning of the first time series. The motion-corrected functional volumes were then coregistered to each participant’s high-resolution T1-weighted anatomical scan using function *spm_coreg* in SPM12. Next, a general linear model was implemented using *3dDeconvolve* in AFNI, including two boxcar regressors, one for left-sided stimuli and one for right-sided stimuli, to estimate condition-specific beta maps.

## Results

### Brain mapping and TUS engagement with the early visual cortex

To confirm accurate targeting and effective engagement of the left early visual cortex, we integrated post-session transcranial acoustic simulations with the functional MRI data. As shown in Figure 2, right hemifield stimulation revealed BOLD activation in the left occipital cortex (Fig.2A), while PETRA-derived pseudo-CT images supported detailed modelling of skull acoustic properties (Fig.2B). The resulting simulations (Fig.2C) indicated focused ultrasound energy delivery to the cortical surface, and critically, the stimulation focus overlapped with the functionally defined target in the left early visual cortex (Fig.2D). Target engagement was quantified as the percentage overlap between the region exceeding 150 kPa in the simulated acoustic field and the BOLD-activated voxels in the functionally defined target region. The average MNI coordinates and standard deviations for the targeted region are × = −10.5 (4.2), Y = −89 (4.0), Z = 3.6mm (5.3).

**Fig. 2.**
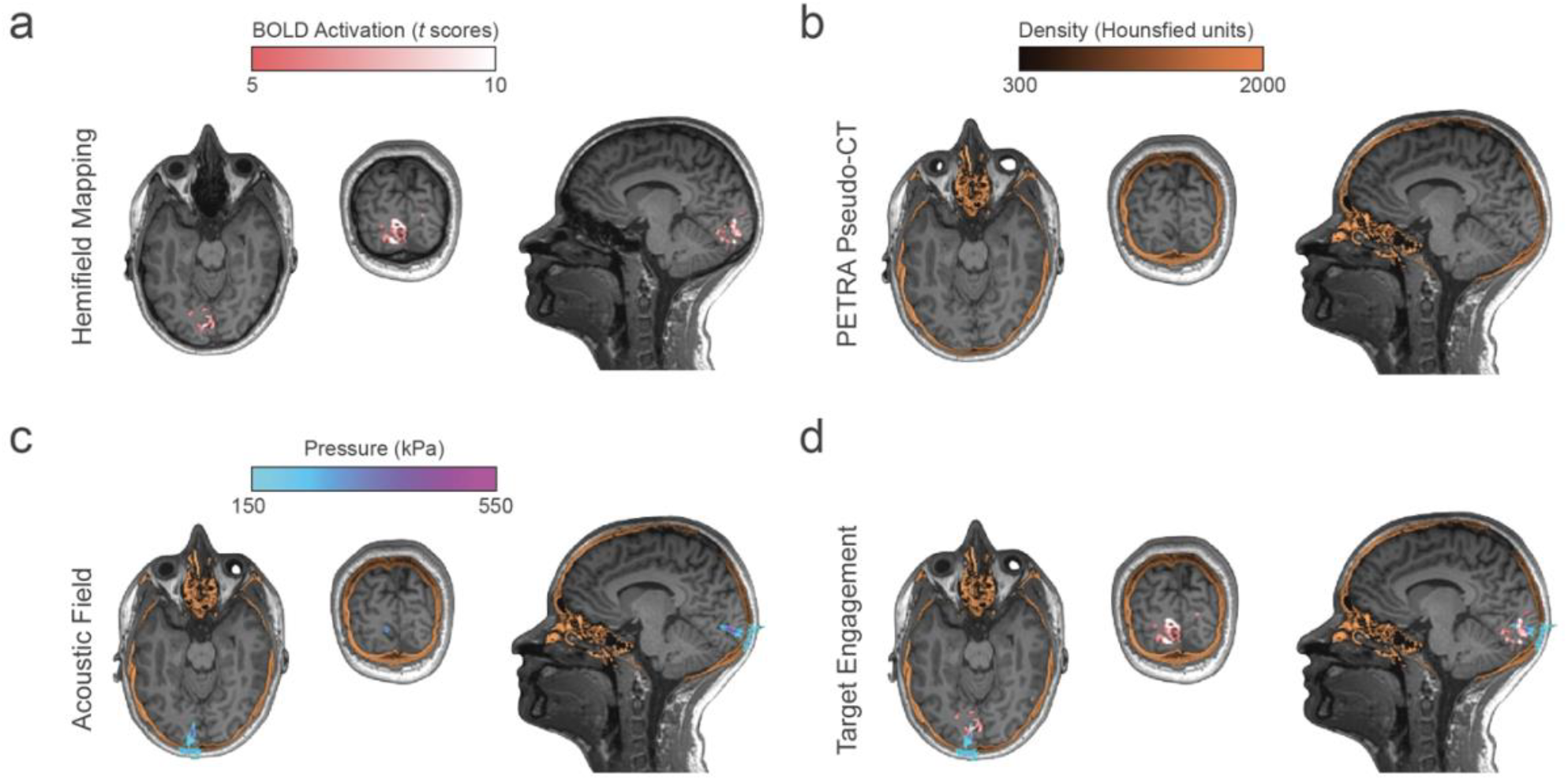
Hemifield mapping and post-session transcranial acoustic simulations in a representative individual. **(A)** T1-weighted MRI image with right hemifield stimulation BOLD response (t-scores). **(B)** Pseudo-CT derived from a PETRA (pointwise encoding time reduction with radial acquisition) MRI sequence, used to estimate skull acoustic properties. (**C)** Simulated ultrasound intensity (pressure) overlaid on the T1-weighted MRI. (**D)** Reliable targeting of the left early visual cortex (upper bank of the left calcarine sulcus, in this example), with target derived from overlaid BOLD response.

Online ultrasound exposure parameters were assessed at the ROI within the left calcarine sulcus target, which used hemifield/quadrant mapping data, as well as in soft tissue and within the skull (Table 1). Temperature increases, measured as thermal dose (CEM43°C), remained minimal across all tissue types, with the highest value in the skull (0.03 ± 0.03 CEM43°C). The transcranial mechanical index (MI) also remained within a safe range across all measurements, with average values of 0.55 ± 0.10 in soft tissue, 0.44 ± 0.13 in the ROI, and 0.73 ± 0.19 in the skull, indicating compliance with established safety limits (43). Pressure levels varied by tissue, with the skull showing the greatest attenuation with the highest intensity spatial peak pulse average (ISPPA) values, ranging from 4.03 to 17.34 W/cm^2^ depending on individual variability. Corresponding pressure values averaged at 0.31 ± 0.09 MPa in the ROI, 0.39 ± 0.07 MPa in the soft tissue, and 0.52 ± 0.13 MPa in the skull. Ultrasound exposure parameters are detailed in Table 1. All measurements remained within established safety limits.

### Behavior

Participants performed the task with high accuracy across all conditions, indicating that task demands were well matched. Response accuracy was consistently high and ranged from 95.8% to 96.3%. A 2×2 repeated-measures ANOVA revealed no significant main effect of stimulation or hemifield, and no interaction between these factors (all *F*s < 0.6, all *p*s > 0.4, all *η*^*2*^*p* < 0.04).

### VEPs

A median of 123 trials were averaged for each condition and participant. Typical pattern onset VEPs were elicited by the visual hemifield stimuli. Since the visual stimuli were large and encompassed both upper and lower visual quadrants (Fig. 3A), the C1 component found in some other studies partially cancelled out and merged with the temporally overlapping P1 component (e.g., 44, 45), making the P1 the first visible peak. This component, peaking around 100 ms, showed a clearly lateralized topography with greater amplitude contralateral to the stimulated hemifield, (Fig. 3C).

**Fig. 3.**
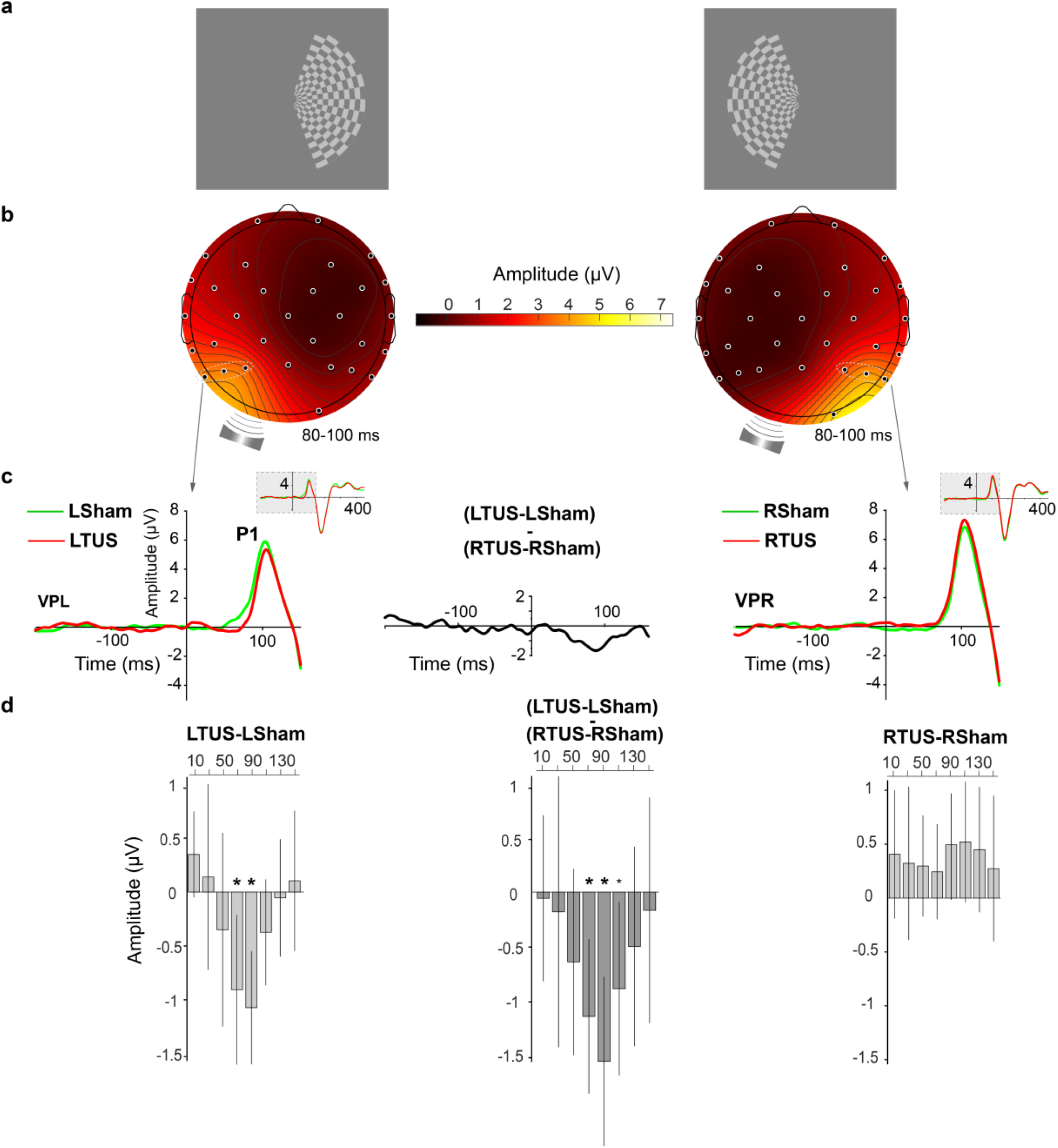
TUS modulation of VEPs. **(A)** Checkerboards (25% contrast) used to stimulate the two visual hemifields. (**B)** Topographic voltage patterns elicited by the contralateral checkerboard in the Sham condition between 80-100 ms, when the effects of TUS were maximal (just before the P1 peak). The TUS transducer was placed over the left occipital cortex. The dashed ellipses highlight electrodes P3, P5, and P7 (left, experimental hemisphere), and P4, P6, and P8 (right, control hemisphere) used to derive virtual electrodes VPL and VPR, respectively. **(C)** Left and right panels show VEPs at VPL and VPR, respectively, during the first 160 ms after contralateral checkerboard stimulation for Sham and TUS trials. The baseline is −200 to 0 ms, and the smaller insets display ERP waveforms from −200 to 450 ms. The central panel (black solid line) shows the time course of the difference between TUS-Sham differences recorded at VPL and VPR (i.e., the interaction). **(D)** Left and right panels show the time course of the voltage difference between TUS and Sham at VPL (LTUS-LSham) and VPR (RTUS-RSham), respectively. The central panel shows the interaction between VPL and VPR ((LTUS-LSham) - (RTUS-RSham)), calculated to eliminate potential effects of sound artifacts. Large asterisks denote time windows surviving multiple comparison correction, while small asterisks indicate time windows significant only before correction (p < .03). All bars are shown with 97% confidence intervals.

The first analysis comparing TUS and sham conditions (LTUS – LSham) at virtual electrode VPL found a significant difference at 60-80 ms, *t*(18) = 3.075, *p* = .007, 97% CI (0.212, 1.602), and at 80-100 ms, *t*(18) = 4.819, *p* = .001, 97% CI (0.548, 1.598), with both effects surviving multiple comparison correction. During this time, preceding the P1 peak, TUS reduced the amplitude of the VEPs (Fig. 3C). This same analysis carried out at control site VPR (RTUS – RSham), showed no significant effects in any time windows (Fig. 3D).

To address the key question of whether the differences at VPL could be due to non-specific effects such as auditory artifacts, we compared them to the corresponding differences computed for virtual electrode VPR, (LTUS - LSham) - (RTUS - RSham). This contrast should eliminate non-specific effects occurring in both hemispheres. Results confirmed the findings of the first analysis and showed a significant VEP reduction between 60 and 120 ms: 60-80 ms, *t*(18) = 3.789, *p* = .001, 97% CI (0.435, 1.867), 80-100 ms, *t*(18) = 4.719, *p* = .0002, 97% CI (0.785, 2.351), and 100-120 ms, *t*(18) = 2.624, *p* = .017, 97% CI (0.091, 1.697). The first two time windows survived multiple comparison correction, highlighting targeted neuromodulatory effects of TUS on early visual processing (Fig. 3D). Finally, to check whether this effect varied by block (e.g., due to practice or carryover effects), we conducted a 3-way ANOVA on the data within the time window with the largest effect (80-100 ms). This ANOVA confirmed the t-test findings, with a main effect of hemisphere, *F*(1,18) = 34.81, *p* < .001, *η*^*2*^*p* = .659 and an interaction between hemisphere and condition, *F*(1,18) = 26.29, *p* < .001, *η*^*2*^*p* = .594. There were no significant effects involving the block factor, suggesting that the effects were consistent across blocks.

### P1 fractional peak latency

A 2 × 2 ANOVA on 50% fractional P1 latency revealed a significant interaction between hemisphere and condition, *F*(1,18) = 6.229, *p* = .023, *η*^*2*^*p* = .257. Follow-up analyses to unpack this interaction showed that P1 50% fractional peak latency at VPL was later for TUS (92ms) than for Sham (86ms), (*t*(18) = 2.18, *p*=.043), suggesting that TUS modulation not only suppressed incoming neural activity, but also slowed it down. In contrast at VPR, P1 fractional peak latency for TUS (90 ms) and Sham (92 ms) did not differ, *t*(18) = −1.39, *p*=.183.

### Target engagement

We defined target engagement from the acoustic simulations as the overlap between the area of stimulation (area of the pressure field above 150 kPa) and the functionally defined target region from retinotopic mapping, quantified as the percentage of BOLD-activated voxels falling within the high-pressure focal zone (Z threshold = 3). A linear mixed-effects analysis revealed a significant positive relationship between Target Engagement (TE) and the VEP difference TUS-Sham between VPL and VPR in the 80-100 ms time window, controlling for experimental blocks and accounting for individual subject variability (Estimate = 0.089, SE = 0.03, *t*(74) = 2.91, *p* =.004). Specifically, higher target engagement corresponded to increased asymmetry between effects at VPL and VPR. As shown in Figure 4, the initial scatter plot, encompassing all data points, showed a modest but statistically significant correlation (one-tailed test: *r* = 0.19, *p* = 0.04). Subsequent scatter plots separated by experimental blocks consistently support this trend, demonstrating the robustness of the relationship across conditions. These findings suggest that greater target engagement is reliably associated with greater differential neural modulation by TUS.

**Fig. 4.**
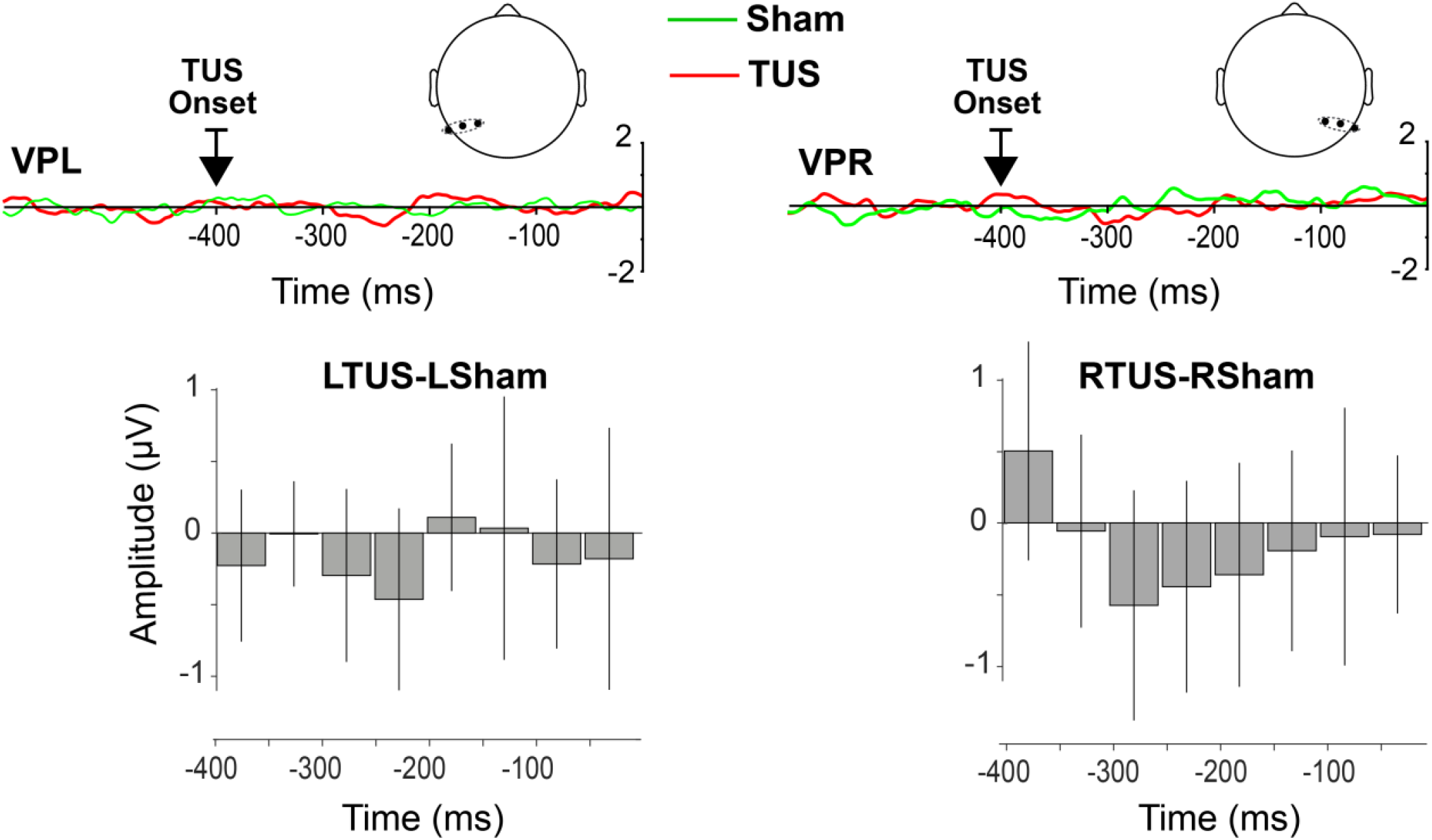
Pre-stimulus EEG. The top panels depict EEG activity time-locked to stimulus onset, shown from −600 to 0 ms at virtual electrodes VPL and VPR. The baseline is from −600 to −400ms, and TUS onset occurs at −400 ms. Importantly, TUS onset itself elicited no VEPs. The bottom panels show the results of sequential t-tests conducted in 50 ms time windows between −400 and 0 ms, comparing the TUS and Sham conditions at VPL and VPR. No significant differences were observed in any of the time windows. Note that comparing VPL and VPR in this case is not meaningful because the EEG data precedes lateralized stimulus onset.

**Fig. 5.**
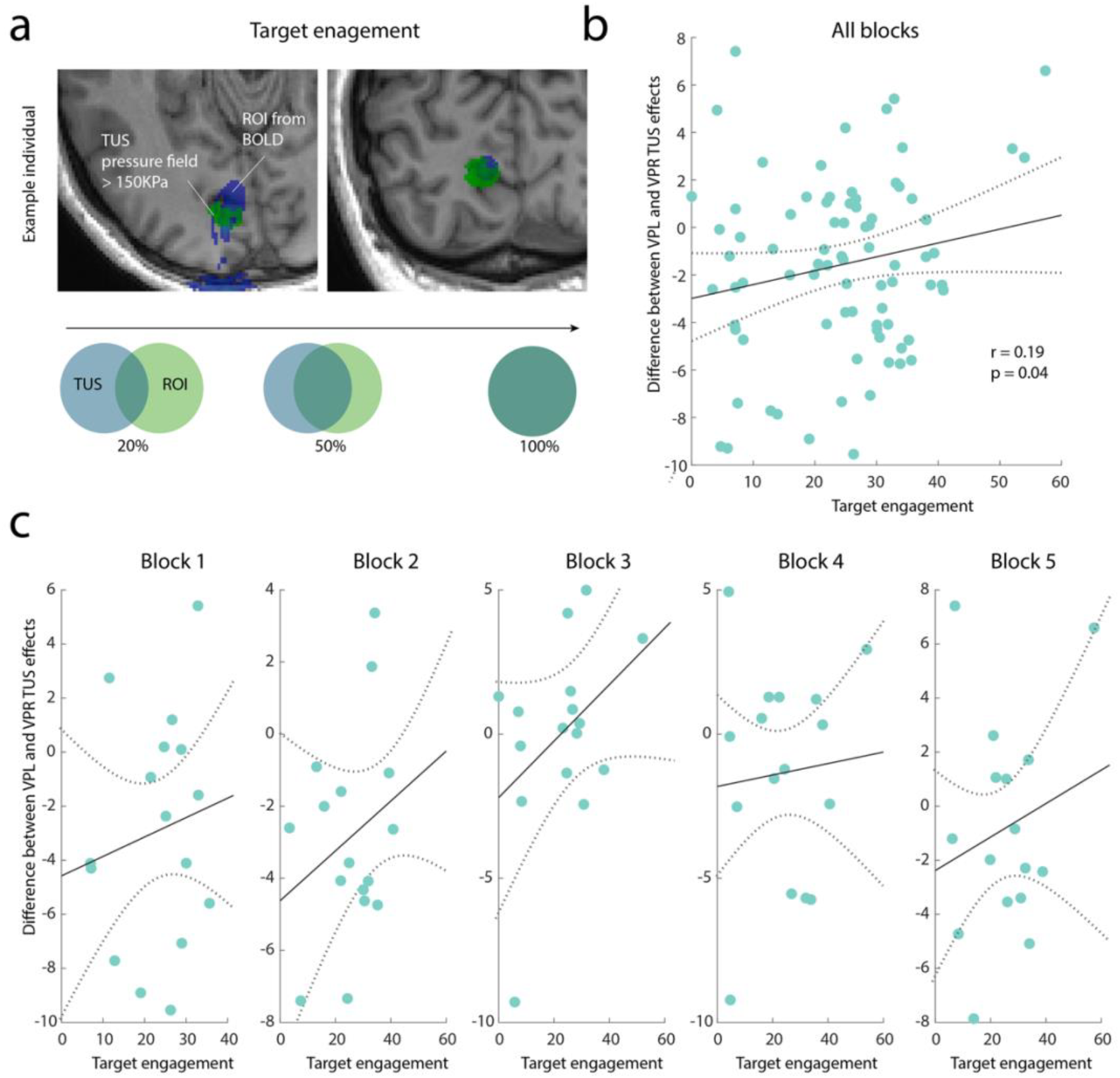
Relationship between target engagement and TUS effects on VEPs. **a** Close-up views of a representative participant showing overlap between simulated ultrasound pressure (blue) and BOLD-defined early visual cortex activation (green). Illustration of the overlap levels used to calculate target engagement, defined as the proportion of BOLD activation within the high-pressure focal zone (>150 kPa). **b** Across all blocks, greater target engagement was associated with larger differential VEP responses (TUS–Sham for contralateral stimuli 80 to 100 ms) between left and right virtual electrodes VPL and VPR (r = 0.19, *p* =.04), indicating that individuals with more precise targeting showed stronger lateralized effects. **c** This relationship was further explored across five separate task blocks. Positive correlations between target engagement and VEP modulation were observed across all blocks, consistently trending in the same direction above and beyond block-wise variability.

## Discussion

Demonstrating online neuromodulatory effects has been an enduring challenge in human TUS neuromodulation research. Here, we provide evidence of such an effect, targeting lateralized responses in early visual cortex. Our design and execution effectively rules out auditory confounds, which have posed a well-recognized challenge in the TUS literature and would have produced bilateral effects. Moreover, we found that greater target engagement, defined as the anatomical overlap between the BOLD-activated voxels and the TUS high-pressure focal zone, was associated with stronger neural modulation. This further supports the validity of our results and underscores the importance of rigorous targeting and simulation. In addition, online paradigms, such as the one employed here, offer methodological advantages over offline approaches, as all conditions are intermixed within the same session. This minimizes confounding effects related to arousal, fatigue, or scanner drift, and enhances the interpretability of observed neuromodulatory changes.

Although the specific pattern of VEPs found in this study differed from that found in Nandi et al (26), some aspects of the findings were similar. Specifically, in both studies TUS modulated the VEPs before 100 ms post-stimulus, and in neither case did TUS alone elicit VEPs or neuromodulation effects. The differences in VEP patterns are not unexpected, as these depend on the spatiotemporal parameters of the visual stimuli. Nandi et al (26) reported neuromodulatory effects on a parietal N75 component that was absent in our data, likely reflecting differences in stimulus paradigms. Specifically, they used a pattern-reversal stimulus (a full-field flickering checkerboard with a 2 Hz reversal rate), which typically elicits an N75 response (e.g., 46), whereas we employed a pattern onset/offset stimulus, which more commonly evokes C1/P1 components (28). Furthermore, the reference site used in their study was not reported, which could also contribute to topographical differences. In our data, the first peak was a large positive component contralateral to the stimulus, consistent with a P1, peaking around 100 ms. We did not observe an earlier C1 component, likely because hemifield stimuli can evoke C1 components of opposite polarity (positive for lower-field stimuli, negative for upper-field stimuli) that partially cancel each other out and merge with the large and temporally overlapping P1 (44, 45).

We observed a reduction in VEP amplitude between 60 and 100 ms after stimulus onset, preceding the P1 peak rather than overlapping with any specific VEP peak. However, the peaks visible in scalp recordings simply represent the summed activity of multiple neural generators in visual cortex, and do not hold any significance over other portions of the VEP slope (47). Peaks recorded at the scalp do not necessarily correspond to discrete neural processes; instead, the relevant information lies in the timing and pattern of differences. The timing and topography of our effects are consistent with, though not limited to, modulation of early visual cortex contralateral to the stimulated visual hemifield. While P1 amplitude was somewhat larger over the right hemisphere (Fig. 2C), consistent with prior reports of hemispheric asymmetry in early responses to large visual stimuli (e.g., 48), this asymmetry did not affect our cross-hemisphere contrast (Fig. 2D), as the key outcome was the difference between TUS and Sham conditions across hemispheres. In addition to a VEP amplitude reduction, TUS also produced a slight increase in the latency of the P1 peak.

Although our intended stimulation target lay within fMRI-activated voxels in the calcarine sulcus, which contains most of V1, the practical limitations of neuronavigation accuracy and transducer stability, even when held in place using a mechanical arm, make it difficult to ensure that stimulation was restricted to V1. Small positional drifts over time were unavoidable, even with anatomy-guided targeting. For this reason, we refer more broadly to early visual cortex rather than V1 specifically. This uncertainty also motivated our use of hemifield stimulation instead of a small visual patch. While a spatially localized stimulus could have engaged a more confined cortical area, it would have carried a higher risk of missing the target entirely due to navigation error. In contrast, modest targeting deviations with stimulation remaining in the left hemisphere, would still affect portions of the early visual cortex responsive to right hemifield input. Although this approach reduced spatial specificity, it provided a pragmatic trade-off under current technical constraints. A potential concern with this approach is that activation of cortical regions outside the focal TUS zone may introduce additional neural signals at the scalp, thereby reducing the relative contribution of modulated activity to the overall EEG signal and decreasing our ability to detect TUS effects. However, the work by Nandi et al. (26) demonstrated early VEP modulation in even less favorable conditions (i.e., full-field stimulation without neuronavigation), supporting the feasibility of our design despite its broader cortical engagement. In future studies, transducer helmets (49) or wearable systems (50) may offer improved mechanical stability could help maintain precise targeting over longer sessions and enable investigation of TUS effects with more spatially constrained stimuli.

Nonetheless, our results show that greater target engagement, defined as the overlap between the high-pressure stimulation focus and functionally activated early visual cortex, was associated with stronger neural modulation effects. While there remains considerable uncertainty in estimating the exact maximum pressure delivered to the cortex from acoustic simulations, target engagement may serve as a more reliable indicator of spatial accuracy (51). This is because it reflects not only intensity but also how precisely the stimulation focus aligns with the intended neural target. These findings suggest that individualised fMRI-guided targeting combined with measures of target engagement can improve the interpretation and efficacy of TUS by ensuring that stimulation reaches the neural populations most relevant to the experimental or clinical goals.

Recent work by Grigoras et al. (49) highlights a promising direction for online TUS research, combining real-time stimulation with concurrent fMRI to study the effects of sonication in the human lateral geniculate nucleus (LGN). Their use of in-scanner targeting allows for precise individualized target engagement, offering a level of accuracy beyond what was achievable in the present study. Furthermore, their custom-built transducer helmet with a significantly higher number of elements (256 elements) enables more precise acoustic focusing than the device used in the current study (4 elements), with dynamic 3D focusing and steering. The LGN is a subcortical relay, so as a target it may offer advantages over cortical regions such as V1, energy deposition in the smaller volume may produce more consistent downstream modulation of early visual processing, potentially leading to more distributed and detectable effects. While many aspects of the experimental design by Grigoras et al. (49) are superior to ours, it is also significantly more resource-intensive, requiring a bespoke transducer system and in-scanner implementation. Nonetheless, it illustrates a potential future direction for online TUS research, where simultaneous targeting and real-time neurofeedback within the MRI environment could serve as a powerful testbed for refining stimulation protocols and improving mechanistic understanding.

## Funding sources

Pump Priming Funds from the School of Psychology and from the Bain Research Imaging Centre, University of Plymouth (SD)

UK Research and Innovation, Future Leaders Fellowship grant MR/Y034368/1 (EF) Biotechnology and Biological Sciences Research Council grant BB/Y001494/1 (EF) Engineering and Physical Sciences Research Council, Neuromod+ Programme grant EP/W035057/1 (EF)

Advanced Research and Invention Agency grant SCNI-PR01-P15 (EF)

### Author contributions

Conceptualization: EF, GG Data curation: SD, GG Formal analysis: SD, EF, GG Funding acquisition: EF, GG Investigation: SD, ED, EF, GG Methodology: SD, KM, EF, GG Project administration: SD Resources: SD, EF, GG Software: SD, EF, GG Supervision: EF, GG Validation: SD, EF, GG Visualization: SD, EF, GG Writing—original draft: SD, EF, GG Writing—review & editing: SD, KM, EF, GG

## Data and materials availability

All data, code, and materials used in the analyses are available from the corresponding author upon reasonable request. All data are available in the main text or the supplementary materials. There are no restrictions on data access.

## References

1. G. Darmani, T. O. Bergmann, K. Butts Pauly, C. F. Caskey, L. de Lecea, A. Fomenko, E. Fouragnan, W. Legon, K. R. Murphy, T. Nandi, M. A. Phipps, G. Pinton, H. Ramezanpour, J. Sallet, S. N. Yaakub, S. S. Yoo, R. Chen, Non-invasive transcranial ultrasound stimulation for neuromodulation. Clin. Neurophysiol. 135, 51–73 (2022).

2. K. Murphy, E. Fouragnan, The future of transcranial ultrasound as a precision brain interface. PLoS Biol. 22, e3002884 (2024).

3. S. N. Yaakub, N. Bault, M. Lojkiewiez, E. Bellec, J. Roberts, N. S. Philip, M. F. S. Rushworth, E. F. Fouragnan, Non-invasive Ultrasound Deep Neuromodulation of the Human Nucleus Accumbens Increases Win-Stay Behaviour. bioRxiv., 2024.2007.2025.605068 (2024).

4. N. Bault, S. N. Yaakub, E. Fouragnan, Early-phase neuroplasticity induced by offline transcranial ultrasound stimulation in primates. Curr. Opin. Behav. Sci. 56, 101370 (2024).

5. B. Clennell, T. G. J. Steward, M. Elley, E. Shin, M. Weston, B. W. Drinkwater, D. J. Whitcomb, Transient ultrasound stimulation has lasting effects on neuronal excitability. Brain Stimul. 14, 217–225 (2021).

6. S. N. Yaakub, T. A. White, J. Roberts, E. Martin, L. Verhagen, C. J. Stagg, S. Hall, E. F. Fouragnan, Transcranial focused ultrasound-mediated neurochemical and functional connectivity changes in deep cortical regions in humans. Nat. Commun. 14, 5318 (2023).

7. T. S. Riis, D. A. Feldman, A. J. Losser, A. Okifuji, J. Kubanek, Noninvasive targeted modulation of pain circuits with focused ultrasonic waves. PAIN. 165, 2829–2839 (2024).

8. C. Aurup, H. A. S. Kamimura, E. E. Konofagou, High-Resolution Focused Ultrasound Neuromodulation Induces Limb-Specific Motor Responses in Mice in Vivo. Ultrasound Med. Biol. 47, 998–1013 (2021).

9. S. S. Yoo, A. Bystritsky, J. H. Lee, Y. Zhang, K. Fischer, B. K. Min, N. J. McDannold, A. Pascual-Leone, F. A. Jolesz, Focused ultrasound modulates region-specific brain activity. Neuroimage. 56, 1267–1275 (2011).

10. K. R. Murphy, J. S. Farrell, J. Bendig, A. Mitra, C. Luff, I. A. Stelzer, H. Yamaguchi, C. C. Angelakos, M. Choi, W. Bian, T. DiIanni, E. M. Pujol, N. Matosevich, R. Airan, B. Gaudillière, E. E. Konofagou, K. Butts-Pauly, I. Soltesz, L. de Lecea, Optimized ultrasound neuromodulation for non-invasive control of behavior and physiology. Neuron. 112, 3252-3266.e3255 (2024).

11. T. Nandi, B. R. Kop, K. Butts Pauly, C. J. Stagg, L. Verhagen, The relationship between parameters and effects in transcranial ultrasonic stimulation. ArXiv., (2024).

12. T. Nandi, B. R. Kop, K. Naftchi-Ardebili, C. J. Stagg, K. B. Pauly, L. Verhagen, Biophysical effects and neuromodulatory dose of transcranial ultrasonic stimulation. Brain Stimul. 18, 659–664 (2025).

13. K. R. Murphy, T. Nandi, B. Kop, T. Osada, M. Lueckel, W. A. N’Djin, K. A. Caulfield, A. Fomenko, H. R. Siebner, Y. Ugawa, L. Verhagen, S. Bestmann, E. Martin, K. Butts Pauly, E. Fouragnan, T. O. Bergmann, A practical guide to transcranial ultrasonic stimulation from the IFCN-endorsed ITRUSST consortium. Clin. Neurophysiol. 171, 192–226 (2025).

14. C. Bimbard, T. P. H. Sit, A. Lebedeva, C. B. Reddy, K. D. Harris, M. Carandini, Behavioral origin of sound-evoked activity in mouse visual cortex. Nat. Neurosci. 26, 251–258 (2023).

15. B. R. Kop, L. de Jong, K. B. Pauly, H. E. M. den Ouden, L. Verhagen, Parameter optimisation for mitigating somatosensory confounds during transcranial ultrasonic stimulation. Brain Stimul., (2025).

16. A. Johnstone, T. Nandi, E. Martin, S. Bestmann, C. Stagg, B. Treeby, A range of pulses commonly used for human transcranial ultrasound stimulation are clearly audible. Brain Stimul. 14, 1353–1355 (2021).

17. L. C. Sincich, J. C. Horton, The circuitry of V1 and V2: integration of color, form, and motion. Annu. Rev. Neurosci. 28, 303–326 (2005).

18. E. A. DeYoe, G. J. Carman, P. Bandettini, S. Glickman, J. Wieser, R. Cox, D. Miller, J. Neitz, Mapping striate and extrastriate visual areas in human cerebral cortex. Proc. Natl. Acad. Sci. U. S. A. 93, 2382–2386 (1996).

19. S. A. Engel, G. H. Glover, B. A. Wandell, Retinotopic organization in human visual cortex and the spatial precision of functional MRI. Cereb. Cortex. 7, 181–192 (1997).

20. M. I. Sereno, A. M. Dale, J. B. Reppas, K. K. Kwong, J. W. Belliveau, T. J. Brady, B. R. Rosen, R. B. Tootell, Borders of multiple visual areas in humans revealed by functional magnetic resonance imaging. Science. 268, 889–893 (1995).

21. T. Kammer, K. Puls, M. Erb, W. Grodd, Transcranial magnetic stimulation in the visual system. II. Characterization of induced phosphenes and scotomas. Exp. Brain. Res. 160, 129–140 (2005).

22. S. Kastner, I. Demmer, U. Ziemann, Transient visual field defects induced by transcranial magnetic stimulation over human occipital pole. Exp. Brain. Res. 118, 19–26 (1998).

23. G. Thut, G. Northoff, J. R. Ives, Y. Kamitani, A. Pfennig, F. Kampmann, D. L. Schomer, Pascual-Leone, Effects of single-pulse transcranial magnetic stimulation (TMS) on functional brain activity: a combined event-related TMS and evoked potential study. Clin. Neurophysiol. 114, 2071–2080 (2003).

24. A. Reichenbach, K. Whittingstall, A. Thielscher, Effects of transcranial magnetic stimulation on visual evoked potentials in a visual suppression task. Neuroimage. 54, 1375–1384 (2011).

25. B. R. Kop, Y. Shamli Oghli, T. C. Grippe, T. Nandi, J. Lefkes, S. W. Meijer, S. Farboud, M. Engels, M. Hamani, M. Null, A. Radetz, U. Hassan, G. Darmani, A. Chetverikov, H. E. M. den Ouden, T. O. Bergmann, R. Chen, L. Verhagen, Auditory confounds can drive online effects of transcranial ultrasonic stimulation in humans. Elife. 12, (2024).

26. T. Nandi, A. Johnstone, E. Martin, C. Zich, R. Cooper, S. Bestmann, T. O. Bergmann, B. Treeby, C. J. Stagg, Ramped V1 transcranial ultrasonic stimulation modulates but does not evoke visual evoked potentials. Brain Stimul. 16, 553–555 (2023).

27. W. Lee, H. C. Kim, Y. Jung, Y. A. Chung, I. U. Song, J. H. Lee, S. S. Yoo, Transcranial focused ultrasound stimulation of human primary visual cortex. Sci Rep 6, 34026 (2016).

28. W. R. Biersdorf, Different scalp localization of pattern onset and reversal visual evoked potentials. Doc. Ophthalmol. 66, 313–320 (1987).

29. S. Yoo, D. R. Mittelstein, R. C. Hurt, J. Lacroix, M. G. Shapiro, Focused ultrasound excites cortical neurons via mechanosensitive calcium accumulation and ion channel amplification. Nat. Commun. 13, 493 (2022).

30. E. Martin, J. Jaros, B. E. Treeby, Experimental Validation of k-Wave: Nonlinear Wave Propagation in Layered, Absorbing Fluid Media. IEEE Trans. Ultrason. Ferroelectr. Freq. Control. 67, 81–91 (2020).

31. J.-F. Aubry, D. Attali, M. Schafer, E. Fouragnan, C. Caskey, R. Chen, G. Darmani, E. J. Bubrick, J. Sallet, C. Butler, C. Stagg, M. Klein-Flügge, S.-S. Yoo, B. Treeby, L. Verhagen, K. B. Pauly, ITRUSST Consensus on Biophysical Safety for Transcranial Ultrasonic Stimulation. arXiv [physics.bio-ph]., (2024).

32. F. A. Duck, Medical and non-medical protection standards for ultrasound and infrasound. Prog. Biophys. Mol. Biol. 93, 176–191 (2007).

33. A. Delorme, S. Makeig, EEGLAB: an open source toolbox for analysis of single-trial EEG dynamics including independent component analysis. J. Neurosci. Methods. 134, 9–21 (2004).

34. J. Lopez-Calderon, S. J. Luck, ERPLAB: an open-source toolbox for the analysis of event-related potentials. Front. Hum. Neurosci. 8, 213 (2014).

35. A. de Cheveigne, I. Nelken, Filters: When, Why, and How (Not) to Use Them. Neuron. 102, 280–293 (2019).

36. N. Bigdely-Shamlo, T. Mullen, C. Kothe, K. M. Su, K. A. Robbins, The PREP pipeline: standardized preprocessing for large-scale EEG analysis. Front. Neuroinform. 9, 16 (2015).

37. L. Pion-Tonachini, K. Kreutz-Delgado, S. Makeig, ICLabel: An automated electroencephalographic independent component classifier, dataset, and website. Neuroimage. 198, 181–197 (2019).

38. D. Yoshor, W. H. Bosking, G. M. Ghose, J. H. Maunsell, Receptive fields in human visual cortex mapped with surface electrodes. Cereb. Cortex. 17, 2293–2302 (2007).

39. J. J. Foxe, G. V. Simpson, Flow of activation from V1 to frontal cortex in humans. A framework for defining “early” visual processing. Exp. Brain. Res. 142, 139–150 (2002).

40. Y. Benjamini, Y. Hochberg, Controlling the False Discovery Rate: A Practical and Powerful Approach to Multiple Testing. J. R. Stat. Soc., Ser B, Stat. Methodol. 57, 289–300 (2018).

41. B. Fischl, FreeSurfer. Neuroimage. 62, 774–781 (2012).

42. R. W. Cox, AFNI: what a long strange trip it’s been. Neuroimage. 62, 743–747 (2012).

43. E. Martin, J. F. Aubry, M. Schafer, L. Verhagen, B. Treeby, K. B. Pauly, ITRUSST consensus on standardised reporting for transcranial ultrasound stimulation. Brain Stimul. 17, 607–615 (2024).

44. F. Di Russo, A. Martinez, M. I. Sereno, S. Pitzalis, S. A. Hillyard, Cortical sources of the early components of the visual evoked potential. Hum. Brain. Mapp. 15, 95–111 (2002).

45. V. P. Clark, S. Fan, S. A. Hillyard, Identification of early visual evoked potential generators by retinotopic and topographic analyses. Hum. Brain. Mapp. 2, 170–187 (1994).

46. J. V. Odom, M. Bach, M. Brigell, G. E. Holder, D. L. McCulloch, A. Mizota, A. P. Tormene, V. International Society for Clinical Electrophysiology of, ISCEV standard for clinical visual evoked potentials: (2016 update). Doc. Ophthalmol. 133, 1–9 (2016).

47. S. J. Luck, An Introduction to the Event-Related Potential Technique, 2nd ed. (MIT Press, Cambridge, MA, 2014).

48. G. Ganis, M. Kutas, An electrophysiological study of scene effects on object identification. Brain Res. Cogn. Brain Res. 16, 123–144 (2003).

49. I. Grigoras, E. Martin, M. Roberts, O. Wright, T. Nandi, S. Rieger, J. Campbell, T. d. Boer, B. Cox, B. Treeby, C. J. Stagg, Ultrasound neuromodulation of the LGN leads to significant activity changes in the ipsilateral visual cortex of healthy humans. Brain Stimul. 18, 387 (2025).

50. J. M. Fan, K. Woodworth, K. R. Murphy, L. Hinkley, J. L. Cohen, J. Yoshimura, I. Choi, G. Tremblay-McGaw, J. Mergenthaler, C. H. Good, P. A. Pellionisz, A. M. Lee, T. Di Ianni, L. P. Sugrue, A. D. Krystal, Thalamic transcranial ultrasound stimulation in treatment resistant depression. Brain Stimul. 17, 1001–1004 (2024).

51. J. F. Aubry, O. Bates, C. Boehm, K. Butts Pauly, D. Christensen, C. Cueto, P. Gelat, L. Guasch, J. Jaros, Y. Jing, R. Jones, N. Li, P. Marty, H. Montanaro, E. Neufeld, S. Pichardo, G. Pinton, A. Pulkkinen, A. Stanziola, A. Thielscher, B. Treeby, E. van ‘t Wout, Benchmark problems for transcranial ultrasound simulation: Intercomparison of compressional wave models. J. Acoust. Soc. Am. 152, 1003 (2022).

