## Supplementary Materials Table S1. for "Unilateral online ultrasound stimulation of early visual cortex suppresses responses to contralateral visual stimuli"

Table S1.

Ultrasound parameters across different target tissues for left early visual cortex, shown separately for each participant.

| ID | Tissue | Thermal Dose (CEM43°C) | Pressure (MPa) | ISPPA  (W/cm2) | Mechanical Index |
| --- | --- | --- | --- | --- | --- |
| sub-1 | ROI mask | 0.00 ± 0.00 | 0.46 ± 0.02 | 7.06 ± 0.71 | 0.65 ± 0.03 |
|  | Soft tissue | 0.01 ± 0.00 | 0.50 ± 0.01 | 8.06 ± 0.28 | 0.70 ± 0.01 |
|  | Skull | 0.01 ± 0.00 | 0.48 ± 0.03 | 7.52 ± 0.87 | 0.68 ± 0.04 |
| sub-2 | ROI mask | 0.00 ± 0.00 | 0.48 ± 0.04 | 7.58 ± 1.39 | 0.68 ± 0.06 |
|  | Soft tissue | 0.01 ± 0.00 | 0.49 ± 0.04 | 7.78 ± 1.17 | 0.69 ± 0.05 |
|  | Skull | 0.02 ± 0.00 | 0.35 ± 0.02 | 4.03 ± 0.39 | 0.49 ± 0.02 |
| sub-3 | ROI mask | 0.00 ± 0.00 | 0.35 ± 0.02 | 4.12 ± 0.42 | 0.50 ± 0.03 |
|  | Soft tissue | 0.01 ± 0.00 | 0.36 ± 0.01 | 4.26 ± 0.35 | 0.51 ± 0.02 |
|  | Skull | 0.02 ± 0.00 | 0.37 ± 0.03 | 4.40 ± 0.66 | 0.52 ± 0.04 |
| sub-4 | ROI mask | 0.00 ± 0.00 | 0.36 ± 0.01 | 4.20 ± 0.16 | 0.50 ± 0.01 |
|  | Soft tissue | 0.01 ± 0.00 | 0.41 ± 0.05 | 5.51 ± 1.29 | 0.58 ± 0.07 |
|  | Skull | 0.03 ± 0.01 | 0.65 ± 0.05 | 14.10 ± 2.07 | 0.93 ± 0.07 |
| sub-5 | ROI mask | 0.00 ± 0.00 | 0.42 ± 0.03 | 5.73 ± 0.68 | 0.59 ± 0.04 |
|  | Soft tissue | 0.01 ± 0.00 | 0.46 ± 0.02 | 6.83 ± 0.73 | 0.64 ± 0.03 |
|  | Skull | 0.03 ± 0.00 | 0.55 ± 0.01 | 9.78 ± 0.38 | 0.77 ± 0.02 |
| sub-6 | ROI mask | 0.00 ± 0.00 | 0.38 ± 0.02 | 4.77 ± 0.56 | 0.54 ± 0.03 |
|  | Soft tissue | 0.01 ± 0.00 | 0.39 ± 0.02 | 4.99 ± 0.51 | 0.55 ± 0.03 |
|  | Skull | 0.02 ± 0.01 | 0.38 ± 0.03 | 4.74 ± 0.78 | 0.54 ± 0.04 |
| sub-7 | ROI mask | 0.00 ± 0.00 | 0.31 ± 0.01 | 3.22 ± 0.24 | 0.44 ± 0.02 |
|  | Soft tissue | 0.01 ± 0.00 | 0.31 ± 0.01 | 3.28 ± 0.28 | 0.44 ± 0.02 |
|  | Skull | 0.03 ± 0.01 | 0.55 ± 0.06 | 10.00 ± 2.05 | 0.78 ± 0.09 |
| sub-8 | ROI mask | 0.00 ± 0.00 | 0.29 ± 0.02 | 2.87 ± 0.35 | 0.42 ± 0.03 |
|  | Soft tissue | 0.04 ± 0.02 | 0.53 ± 0.02 | 9.82 ± 0.88 | 0.74 ± 0.03 |
|  | Skull | 0.12 ± 0.09 | 0.72 ± 0.08 | 17.34 ± 4.01 | 1.02 ± 0.12 |
| sub-9 | ROI mask | 0.00 ± 0.00 | 0.12 ± 0.00 | 0.45 ± 0.00 | 0.17 ± 0.00 |
|  | Soft tissue | 0.05 ± 0.00 | 0.41 ± 0.00 | 6.18 ± 0.00 | 0.58 ± 0.00 |
|  | Skull | 0.09 ± 0.00 | 0.58 ± 0.00 | 11.07 ± 0.00 | 0.82 ± 0.00 |
| sub-10 | ROI mask | 0.00 ± 0.00 | 0.35 ± 0.04 | 3.98 ± 0.94 | 0.49 ± 0.06 |
|  | Soft tissue | 0.01 ± 0.00 | 0.36 ± 0.02 | 4.26 ± 0.56 | 0.51 ± 0.03 |
|  | Skull | 0.02 ± 0.00 | 0.55 ± 0.02 | 9.79 ± 0.67 | 0.77 ± 0.03 |
| sub-11 | ROI mask | 0.00 ± 0.00 | 0.22 ± 0.02 | 1.56 ± 0.33 | 0.31 ± 0.03 |
|  | Soft tissue | 0.01 ± 0.00 | 0.36 ± 0.05 | 4.63 ± 0.98 | 0.51 ± 0.06 |
|  | Skull | 0.04 ± 0.01 | 0.71 ± 0.03 | 16.30 ± 1.31 | 1.00 ± 0.04 |
| sub-12 | ROI mask | 0.00 ± 0.00 | 0.25 ± 0.03 | 2.14 ± 0.43 | 0.36 ± 0.04 |
|  | Soft tissue | 0.00 ± 0.00 | 0.44 ± 0.01 | 6.42 ± 0.23 | 0.63 ± 0.01 |
|  | Skull | 0.01 ± 0.00 | 0.37 ± 0.02 | 4.43 ± 0.48 | 0.52 ± 0.03 |
| sub-13 | ROI mask | 0.00 ± 0.00 | 0.33 ± 0.02 | 3.51 ± 0.48 | 0.46 ± 0.03 |
|  | Soft tissue | 0.00 ± 0.00 | 0.37 ± 0.01 | 4.42 ± 0.35 | 0.52 ± 0.02 |
|  | Skull | 0.01 ± 0.00 | 0.41 ± 0.03 | 5.50 ± 0.79 | 0.58 ± 0.04 |
| sub-14 | ROI mask | 0.00 ± 0.00 | 0.34 ± 0.04 | 3.96 ± 0.85 | 0.49 ± 0.05 |
|  | Soft tissue | 0.01 ± 0.00 | 0.38 ± 0.01 | 4.82 ± 0.24 | 0.54 ± 0.02 |
|  | Skull | 0.04 ± 0.01 | 0.63 ± 0.05 | 12.93 ± 2.15 | 0.89 ± 0.07 |
| sub-15 | ROI mask | 0.00 ± 0.00 | 0.25 ± 0.07 | 2.22 ± 0.94 | 0.36 ± 0.10 |
|  | Soft tissue | 0.01 ± 0.00 | 0.30 ± 0.03 | 3.14 ± 0.71 | 0.43 ± 0.04 |
|  | Skull | 0.02 ± 0.00 | 0.51 ± 0.07 | 8.65 ± 2.56 | 0.72 ± 0.10 |
| sub-16 | ROI mask | 0.00 ± 0.00 | 0.22 ± 0.01 | 1.63 ± 0.18 | 0.31 ± 0.02 |
|  | Soft tissue | 0.01 ± 0.00 | 0.31 ± 0.04 | 3.23 ± 0.71 | 0.44 ± 0.05 |
|  | Skull | 0.01 ± 0.00 | 0.39 ± 0.02 | 4.85 ± 0.44 | 0.54 ± 0.03 |
| sub-17 | ROI mask | 0.00 ± 0.00 | 0.27 ± 0.03 | 2.49 ± 0.58 | 0.39 ± 0.04 |
|  | Soft tissue | 0.01 ± 0.00 | 0.35 ± 0.01 | 4.03 ± 0.12 | 0.49 ± 0.01 |
|  | Skull | 0.05 ± 0.01 | 0.70 ± 0.08 | 16.28 ± 3.67 | 0.99 ± 0.11 |

Values for online protocols are given as the mean ± standard deviation for all 5 blocks of the task. Data for 2 participants were not recorded.
